# K_ATP_ channel mutation disrupts hippocampal network activity and nocturnal γ shifts

**DOI:** 10.1101/2023.06.18.545438

**Authors:** Marie-Elisabeth Burkart, Josephine Kurzke, Jorge Vera, Frances M. Ashcroft, Jens Eilers, Kristina Lippmann

**Author notes:** These authors contributed equally (Marie-Elisabeth Burkart, Josephine Kurzke).

## Abstract

ATP-sensitive potassium (K_ATP_) channels enable ATP to control the membrane potential and insulin secretion. Humans affected by severe activating mutations in K_ATP_ channels suffer from developmental *d*elay, *e*pilepsy and *n*eonatal *d*iabetes (DEND syndrome). While the diabetes in DEND syndrome is well understood, the pathophysiology of the neurological symptoms remains unclear. We hypothesized that parvalbumin-positive interneurons (PV-INs) are key for the pathophysiology and found, by using electrophysiology, that expressing the DEND mutation K_ir_6.2-V59M selectively in PV-INs reduced intrinsic gamma frequency preference and short-term depression as well as disturbed cognition-associated gamma oscillations and hippocampal sharp waves. Furthermore, risk of seizures is increased and day-night shift in gamma activity disrupted. Thus, PV-INs play a key role in DEND syndrome and this provides a framework for establishing treatment options.

**One Sentence Summary:** Overactive K_ATP_ channels in PV-interneurons disturb cellular behaviour and cognition-associated network oscillations.

## Main Text

DEND syndrome is a channelopathy caused by activating mutations in either the pore-forming (K_ir_6.2) or regulatory (SUR1) subunits of the K_ATP_ channel (*1–3*) that prevent its inhibition by ATP. Due to the widespread expression of these channels, both endocrine and neurological symptoms prevail. In pancreatic β cells, mutated K_ATP_ channels lead to membrane hyperpolarization and thereby reduce glucose-stimulated electrical activity, calcium influx and insulin release, causing neonatal diabetes (*2–5*). Blockers of K_ATP_ channels, such as the sulfonylureas glibenclamide and tolbutamide, restore insulin secretion and are effective in treating the diabetes of many patients with DEND syndrome (*2, 6, 7*). In contrast, the pathophysiology of the devastating neurological symptoms (developmental delay, seizures, cognitive deficits (*1, 8*– *10*)) remains poorly understood. They result most likely from K_ATP_ channel dysfunction in the CNS (*11, 12*) but it is yet unclear which neurons are involved. Presumably due to unfavorable CNS pharmacokinetics of the drugs (*13, 14*), or mutation-induced drug insensitivities (*2*), the neurological symptoms are largely resistant to K_ATP_ channel blockers. They therefore represent a major challenge in treating DEND syndrome (*8*), demanding a deeper understanding of the pathophysiology at the cellular and network level.

### Open K_ATP_ channels disrupt network activity

We first explored possible network phenomena associated with activating K_ATP_ channel mutations by testing the effects of the K_ATP_ channel opener diazoxide on acute hippocampal slices prepared from wild-type mice. Hippocampal sharp waves (SPWs) and gamma oscillations, known to be relevant for cognitive functions, such as memory consolidation and memory replay as well as for information selection, processing, transfer and learning (*15, 16*), were recorded in the area cornu ammonis 1 (CA1, Fig. 1A). Opening K_ATP_ channels with bath-applied diazoxide (300 μM) halved the number of spontaneously occurring SPWs within 10 minutes, from 77 [55] min^-1^ (median and interquartile range) to a plateau of 34 [28] SPWs min^-1^ (KS test, p=0.0010, Fig. 1B-D) and diminished the SPW amplitude from 0.14 [0.14] mV to 0.09 [0.14] mV (KS test, p=0.0018; Fig. 1E). Moreover, diazoxide reduced the peak frequency of kainate-induced network oscillations from a gamma frequency of 40.3 [3.7] Hz in controls to below gamma (28.1 [25.0] Hz, WSRT p=9.5e-7, Fig. 1F-G). The relative gamma power declined from 80.6 [13.1] % in controls to 52.8 [27.0] % in the presence of diazoxide (paired t-test, p=7.4e-7, Fig. 1H). These findings show that activation of K_ATP_ channels disturbs the generation of distinct patterns of hippocampal oscillatory activity (SPWs and gamma oscillations), which is likely to result in an impairment of cognitive function.

**Fig. 1.**
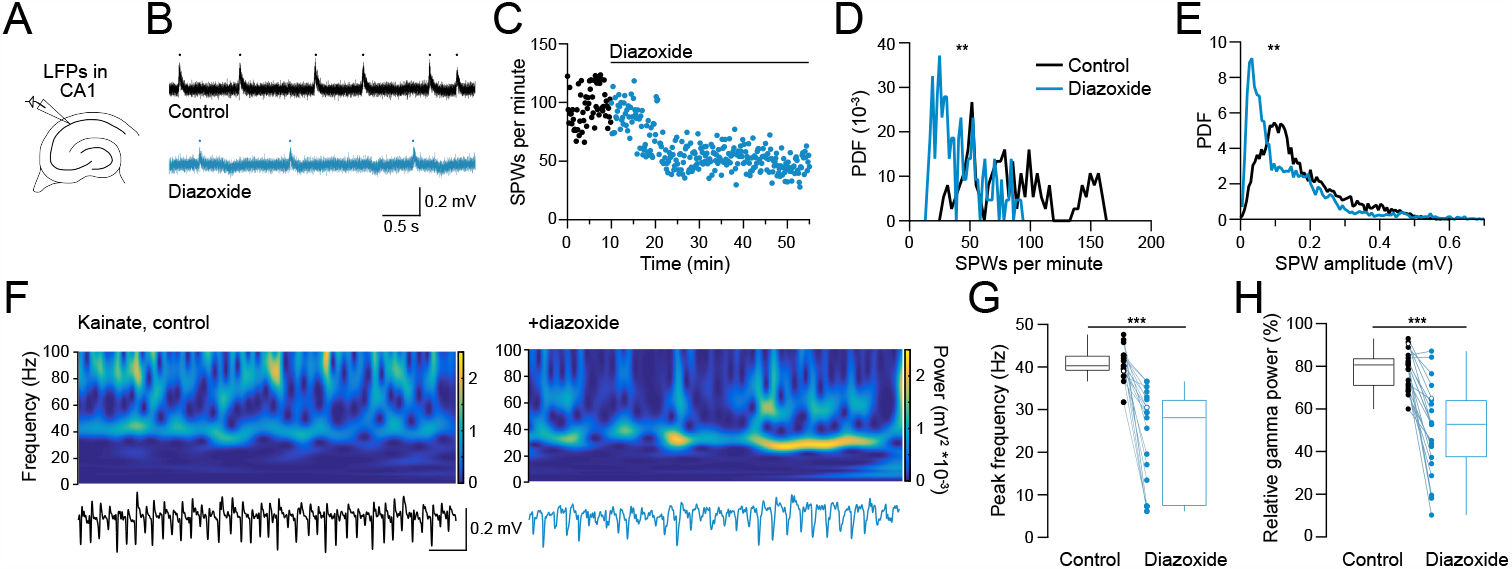
Opening K_ATP_ channels pharmacologically disrupts sharp waves (SPWs) and gamma oscillations in acute hippocampal slices from wild-type mice. **A**. Schematic of recording local field potentials (LFPs) in CA1 of acute hippocampal slices. **B**. Representative LFP recording from a slice in control conditions and in the presence of the K_ATP_ channel opener diazoxide (300 μM; black and blue trace, respectively). SPWs marked by dots. **C**. Corresponding plot of SPWs per min (10 s bins) vs. time before (black symbols) and during wash in of diazoxide (blue symbols). **D**. Corresponding probability density function (PDF) of the group analysis. Data from 10 min periods with 1 min binning. Diazoxide data taken 35 to 45 min after the start of the wash in. Asterisks denote a significant difference (KS, p=0.001). **E**. Corresponding PDF of SPW amplitudes from control (black) and diazoxide (blue) slices (KS, p=0.0018). **F**. Example spectrograms (power spectral densities, PSDs, over time; top) and corresponding LFP recordings (bottom, 200 Hz low-pass filtered) from a slice in which gamma oscillations were induced by prolonged application of 200 nM kainate before (90 min, left) and 45 min after wash-in of diazoxide (in the continued presence of kainate, right). **G-H**. Peak frequency (G) and relative gamma power (H; power from 30-100 Hz relative to the total power from 0.5-100 Hz) before and after diazoxide application. Data were calculated from PSDs covering 5 min periods immediately before and 40 to 45 min after diazoxide application. Box plots and individual data points are shown. Open data circles indicate examples shown in F. Asterisks denote significant differences in the peak frequency (WSRT, p=9.5e-7) and relative gamma power (paired t-test, p=7.4e-7).

### A DEND mutation disturbs network activity

The widespread CNS distribution of K_ATP_ channel subunits (K_ir_6.2 and regulatory SUR1, SUR2A and SUR2B subunits (*17–20*)) makes it difficult to predict which brain regions and cell types underlie the various DEND symptoms. It can be envisaged that K_ATP_ channels will be especially relevant for neurons engaged in energy demanding activity like burst firing (*21*), such as neurons involved in generating cognition-associated network activity, whose dysfunction results in epilepsy (*22–24*). For these cells, an activity-induced drop in intracellular ATP and the resulting hyperpolarization due to enhanced K_ATP_ channel activity may represent a feedback mechanism that modulates burst activity and related network phenomena. Furthermore, K_ATP_ channels may also open during metabolic stress to protect neurons from excessive energy loss (*25*).

Previously, it was suggested that inhibitory rather than excitatory hippocampal neurons are equipped with K_ATP_ channels (*26*). We hypothesized that parvalbumin-positive inhibitory interneurons (PV-INs) are likely to underlie the neurological symptoms of DEND syndrome. These fast-spiking cells are essential for the generation of high-frequency, cognition-associated network activity such as SPWs and gamma oscillations (*21, 24, 27, 28*). PV-INs are known to have a high energy-expenditure (*22, 29*). Activity-induced channel opening, metabolic stress, or activating K_ATP_ channel mutations might impair firing of these neurons, the network phenomena they are engaged in, and the inhibitory tone they exert to prevent epileptic activity (*25, 30*).

Therefore, we hypothesized that selectively expressing a DEND mutation in PV-INs should lead to similar network effects as those observed following bath application of diazoxide. To this end, using a PV-Cre reporter line that allows targeting of PV-INs (Fig. 2A), we created a mutant mouse (PV-V59M) expressing the activating mutation K_ir_6.2-Val59 -> Met59 (the most common human DEND mutation (*2, 12*)) selectively in PV-INs. And, indeed, mimicking the effects of diazoxide, hippocampal slices from PV-V59M mice showed a significantly reduced rate of spontaneous SPWs (38 [72] min^-1^ vs. 64 [70] min^-1^ in littermate control mice (denoted as ‘littermates’ throughout the figures and text, see *Methods*), KS test, p=0.0089, Fig. 2B, C) as well as significantly reduced SPW amplitudes (0.04 [0.05] mV vs. 0.07 [0.09] mV in littermates, KS test, p=0.046, Fig. 2D).

**Fig. 2.**
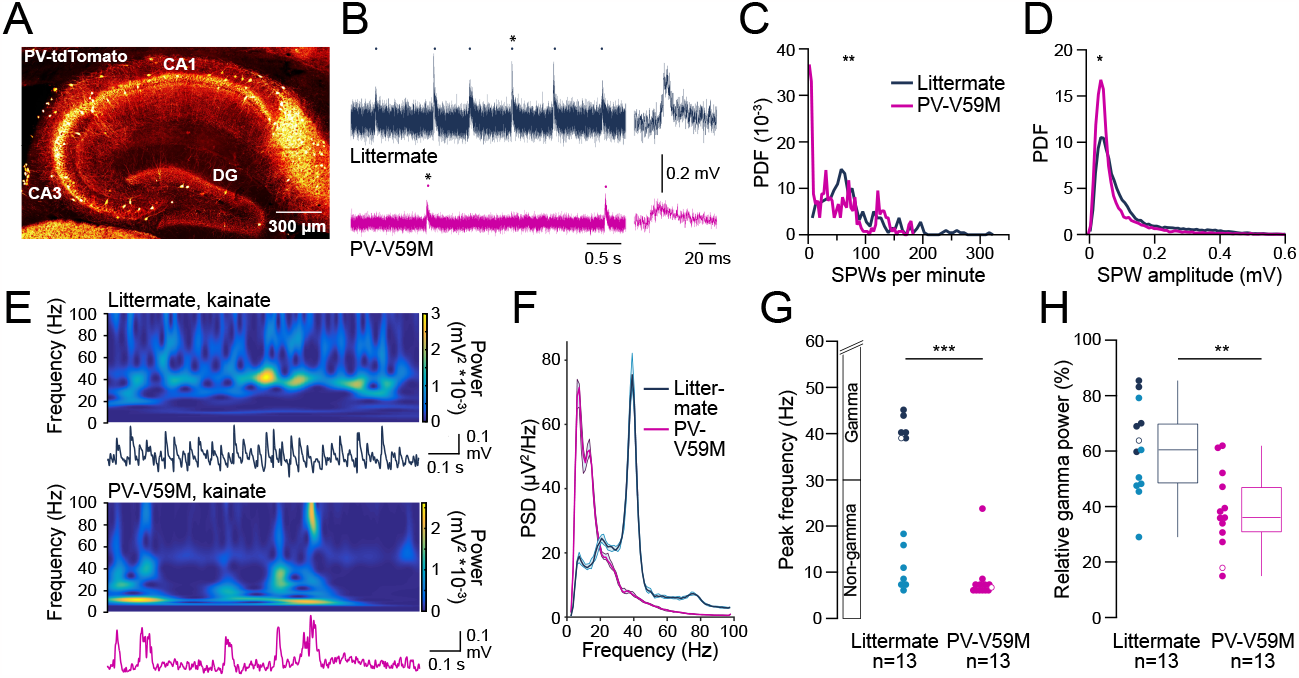
Sharp waves and gamma oscillations are impaired in acute hippocampal slices from PV-V59M mice. **A**. Stitched confocal image of a hippocampal slice from a PV-tdTomato mouse stained with anti-RFP-antibodies (30 μm z-projection taken at 1.5 μm z-interval). PV-INs are indicated in yellow. **B**. Representative LFP recordings from slices from a littermate (dark blue) and a PV-V59M mouse (magenta). Dots mark SPWs. Stars indicate SPWs shown on the right on an expanded time scale. **C-D**. PDFs of SPW frequency (C, 1 min binning) and amplitude (D) in slices from littermates (blue) and PV-V59M mice (magenta). Asterisks denote significant differences (KS test, p=0.0089 in C and 0.046 in D, respectively). **E**. Example spectrograms and corresponding LFP recordings from kainate-treated slices from a littermate (top) and a PV-V59M mouse (bottom). **F**. Corresponding PSDs, computed from the last 5-min data segments after 90 min wash in of kainate. **G-H**. Peak frequency (G) and gamma power (30-100 Hz) relative to the full (0.5-100 Hz) power (H) of recordings from littermate (dark blue for slices having their peak frequency in the gamma range, light blue for the non-gamma range) and PV-V59M slices (magenta). Open circles indicate the examples shown in E-F. Asterisks denote significant differences (MW, p=0.0007 in G and t-test, p=0.001 in H; n denotes the number of slices).

Furthermore, and again mimicking the effects of diazoxide, PV-V59M slices did not generate as strong gamma oscillations as their littermates. About half of the littermate slices (46%, 6 out of 13, mean peak frequency of 18.3 [31.7] Hz) but none of the PV-V59M slices (n=13) showed a peak frequency in the gamma range (mean frequency 6.7 [1.2] Hz, MW test, p=0.0007, Fig. 2 E-G). The relative gamma power in PV-V59M slices was only 36.0 [16.5] % compared to 60.4 [21.9] % in littermates (t-test, p=0.001, Fig. 2H). There was a striking similarity between the effect of the channel opener diazoxide (which affects all neurons that endogenously express K_ATP_ channels) in wild-type slices (Fig. 1) and the effects of the activating K_ATP_ channel mutation in PV-V59M slices (Fig. 2). This suggests (i) that a substantial percentage of K_ATP_ channels are closed in control neurons and (ii) that PV-INs, as key neurons for the generation of SPWs and gamma oscillations, are the predominant neuronal hippocampal cell type affected in DEND syndrome.

### A DEND mutation disturbs cellular activity

In PV-INs the K_ATP_ channel V59M mutation may exert its detrimental effects by altering the intrinsic electrophysiological properties (*31*) of the dendro-somatic compartment, and/or by affecting synaptic release onto postsynaptic targets. Thus, we carried out patch-clamp recordings from single PV-INs. Basic properties (membrane resistance, resting membrane potential, action potential threshold and maximum firing frequency) were unaffected by the V59M mutation (fig. S1A-D). However, the mutation affected two more subtle properties relevant for gamma activity: intrinsic membrane potential oscillations (Fig. 3A-E) and gamma resonance (Fig. 3F-H). Intrinsic oscillatory activity (quantified during long depolarizing voltage steps in periods without action potential firing, Fig. 3A left, cf. (*32*)) revealed patterns that peaked in the gamma range or below (Fig. 3A, right). In littermate neurons, 50% (5 out of 10) of PV-INs peaked in the gamma range and 50% below (‘gamma’ and ‘non-gamma’, respectively), resulting in an average peak frequency of 19 [44] Hz. In contrast, all (n=15) of the PV-V59M neurons showed a peak frequency below gamma at around 2 [1] Hz (MW test, p=0.008 for peak frequency in Fig. 3B and Fisher’s exact test, p=0.005 for categorical differences in Fig. 3C). A power spectral densities (PSDs) plot of the intrinsic oscillations revealed that, in contrast to littermate ‘gamma’ and ‘non-gamma’ neurons, most of the PV-V59M neurons were not able to generate a power plateau in the gamma-band (30-100 Hz, Fig. 3D). Over the gamma frequency range, the power of the intrinsic oscillations was significantly reduced in PV-V59M mice compared to littermates (0.11 [0.06] mV^2^ vs. 0.13 [0.03] mV^2^, MW test, p=0.044, Fig. 3E).

**Fig. 3.**
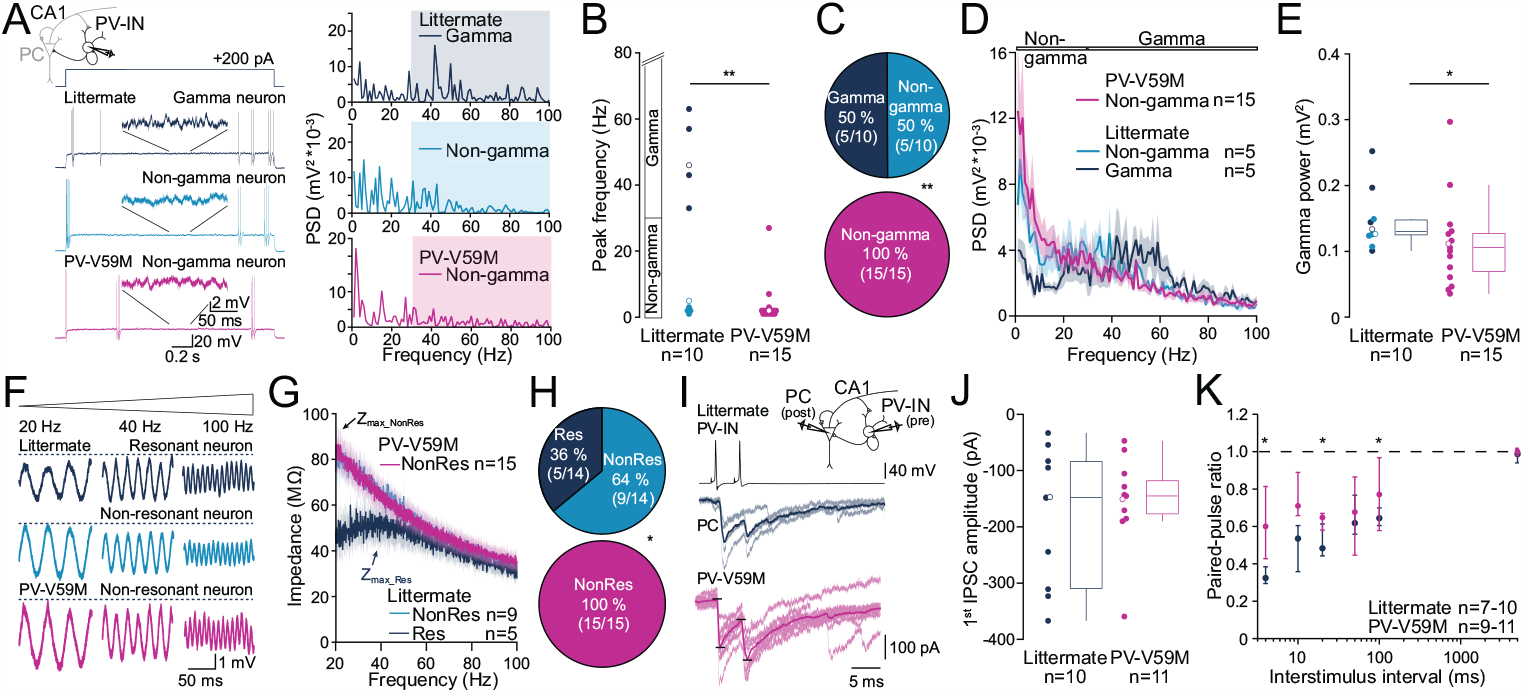
Reduced intrinsic gamma oscillations, gamma resonance and short-term depression in PV-INs of PV-V59M mice. **A**. Left: Examples of intrinsic membrane potential oscillations of a littermate ‘gamma’ (dark blue), ‘non-gamma’ (light blue) and PV-V59M ‘non-gamma’ PV-IN (magenta; color code also applies to B-E). Zoom-ins of the oscillatory activity (upper scale bars). Top: schematic of patch-clamp recording in CA1 PV-INs. Right: Corresponding PSDs obtained from 1 s oscillatory activity without APs. Note: the littermate ‘gamma’ neuron peaked in the gamma band (shaded), whereas the littermate ‘non-gamma’ and PV-V59M neuron peaked in the delta-to-theta range (0.5-8 Hz). **B**. Peak frequency of the PSDs of littermate and PV-V59M PV-INs (MW, p=0.008). Open circles indicate examples from A. **C**. PV-IN proportions with their peak frequency within or below gamma (non-gamma; Fisher’s, p=0.005). **D**. Average PSDs (±SEMs) from PV-INs. **E**. Gamma power of littermate and PV-V59M PV-INs (MW, p=0.044) computed by the area under the curve for the PSDs between 30 and 100 Hz. Open circles denote examples from A. **F**. Example membrane potentials of neurons that responded to a perithreshold ZAP current stimulus (20-100 Hz) either with a resonance peak in the gamma range (top, dark blue, littermate) or not (light blue, littermate; magenta, PV-V59M; same colors for G-H). Zoom-ins at 20, 40 and 100 Hz stimulus frequencies. **G**. Average impedance (±SEM) plotted vs. frequency. Arrows point at the maximal impedance (Z_max_); n=number of neurons. **H**. Proportion of resonant and non-resonant PV-INs from littermate and PV-V59M mice (Fisher’s, p=0.017). **I**. Paired IPSCs evoked in PCs (middle, blue trace: littermate; bottom, magenta trace: PV-V59M) by inducing APs in synaptically connected PV-INs (top). Black lines depict the IPSC quantification. Inset (top) illustrates the recording configuration. **J**. Mean first IPSC amplitudes in PCs. **K**. Paired-pulse ratio of IPSCs (MW, asterisks denote significances (p<0.02) between genotypes).

Gamma resonance at depolarized subthreshold membrane potential (*33*) was tested by applying subthreshold ZAP (impedance amplitude profile) stimuli covering frequencies up to 100 Hz to obtain impedance curves (Fig. 3F). The resonance strength parameter Q was determined from the fitted impedance curves (Fig. 3G; Q=Z_max_fit_/Z_20Hz_fit_, cf. (*34*)) to distinguish between ‘resonant’ (Res, Q>1.05) and ‘non-resonant’ neurons (NonRes, Q ≤ 1.05). Based on this criterion, 36% of PV-INs from control littermates (5 out of 14) resonated in the gamma frequency range, showing their maximum impedance at gamma with 36.5 [3.2] Hz (Fig. 3G). Not a single V59M PV-IN (n=15) revealed gamma resonance behavior (Fig. 3G, magenta trace), revealing a significant categorical difference (Fisher’s, p=0.017, Fig. 3H) between mutant and littermate PV-INs.

Presynaptic effects of the V59M mutation were studied in paired patch-clamp recordings between PV-INs and postsynaptic pyramidal cells (PCs, Fig. 3I, inset), a synaptic connection associated predominantly with the generation of SPWs and gamma oscillations (*27*). Pairs of action potentials evoked in PV-INs reliably evoked pairs of inhibitory post-synaptic currents (IPSCs) in PCs (Fig. 3I). Although the amplitudes of the first IPSCs were similar between mutant and control pairs (Fig. 3I, J), we found a significantly reduced paired-pulse depression (PPD) in PV-V59M INs (Fig. 3I, K; MW test, *p<0.05). A reduction in PPD may result from a variety of pre- and postsynaptic effects (*35*), including complex homeostatic effects (*36*). Nonetheless, functionally, a reduced PPD favors tonic versus phasic synaptic transmission (*35*). Thus, both, the reduced gamma preference (Fig. 3A-H) as well as the reduced PPD (Fig. 3I-K) of PV-V59M INs, potentially contribute to the disturbed network phenotype observed in slices from PV-V59M hippocampi (Fig. 2). This reflects the essential role of PV-INs in the generation of SPWs and gamma oscillations.

### Seizures and absent nocturnal gamma

*In vitro* physiology provides important information regarding the pathophysiology of channelopathies (*37*). However, to determine if PV-INs play a dominant role in the neurological symptoms of patients with DEND syndrome, namely epilepsy and developmental delay (*10, 38*), *in vivo* experiments are required. To this end, we performed chronic (7 days) local field potential (LFP) recordings from the hippocampal area CA1 in freely moving mice (Fig. 4A). Indeed, we found that, while none (n=8) of the littermate mice showed epileptic activity, 7 out of 9 of PV-V59M mice presented with electrographic seizures (Fig. 4B-C, Fisher’s, p=0.002), comprising on average 2.7 [4.3] seizures per day in mutants with a median duration of 41 [32] s per seizure (Fig. 4C-D). Besides epileptic activity, the chronic LFP recordings revealed an unexpected incapacity of PV-V59M mice to adapt their brain activity to the day-night rhythm. Littermates displayed a circadian rhythm characterized electrographically by a pronounced increase in relative gamma power during the night (Fig. 4E-F), which is the time of their wakefulness and activity (day: 28.6 [14.4] % vs. night: 32.9 [18.3] %, paired t-test, p=2e-4, Fig. 4G and left panel in H). While PV-V59M mice showed a normal day-night rhythm in their behavioural activity (fig. S2), they failed to produce a nocturnal increase in gamma power (day: 22.8 [7.6] % vs. night: 23.9 [7.7] %, paired t-test, p=0.379, Fig. 4E-G and right panel in H). Thus, the impaired gamma activity observed in slices of PV-V59M mice is mirrored by an impaired shift in circadian gamma activity *in vivo*, which may affect the ability to select and process information and thus relate to the developmental delay of patients with DEND syndrome.

**Fig. 4.**
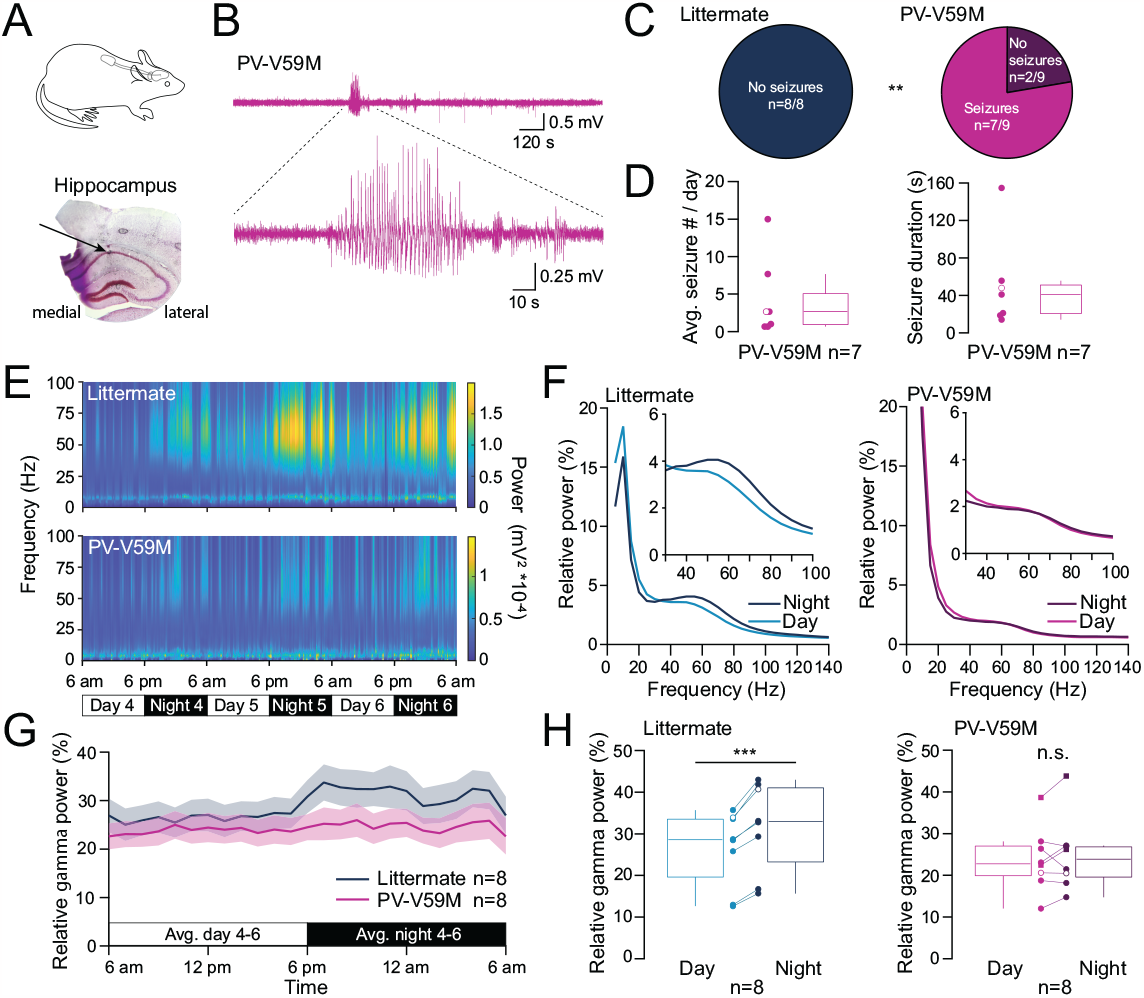
Epileptic seizures and absence of nocturnal increase in hippocampal gamma power in PV-V59M mice. **A**. Top: Schematic of the telemetric hippocampal LFP recording in freely moving mice carrying a transmitter implanted subcutaneously on their back. Bottom: Typical cresyl violet stained hippocampal slice with previously implanted electrode. Arrow points at the recording site in stratum pyramidale of CA1. **B**. Representative recording of *in vivo* hippocampal electrographical activity with a zoom-in of a seizure in a PV-V59M mouse. **C**. Fraction of animals showing electrographical seizures among littermates and PV-V59M mice during postsurgical days 4-6 (Fisher’s, p=0.002). **D**. Average number of seizures per day (left) and median duration of detected seizures (right) in PV-V59M mice. Example from B is indicated with open circles. **E**. Example spectrograms (0.5 Hz binning) of 3-day-long (postsurgical day 4-6) hippocampal LFP recordings from a littermate and a PV-V59M mouse. Note the circadian changes in gamma power in the littermate, which are almost absent in the PV-V59M mouse. Spectrograms are scaled according to their respective maximal power values. **F**. Relative power at night (dark color) and day (light color) plotted vs. frequency for the examples shown in E (littermates: left, dark blue traces; PV-V59M: right, magenta traces, 5 Hz binning). Insets display zoom-ins in the gamma frequency range. **G**. Gamma power over 24 h, averaged over 3 days for 8 littermates and 8 PV-V59M mice, respectively (mean ± SEM). **H**. Relative gamma power at day and night time for littermates (left, paired t-test, p=2e-4) and PV-V59M animals (right, p=0.379). Examples from E-F are depicted with open circles and the two PV-V59M animals without seizures with squares.

## Discussion

These results allow formulation of a refined hypothesis for the neuronal mechanisms underlying the neurological symptoms of DEND syndrome. This proposes that K_ATP_ channels normally regulate the rhythmic firing pattern of neurons that are engaged in burst firing and/or high-frequency network oscillations. Such activity can be expected to lead to a decrease of cytosolic ATP, which, in turn, activates K_ATP_ channels and hyperpolarizes the neurons, ultimately serving as feedback in network oscillations (*26, 29*). Activating K_ATP_ channel mutations such as Kir6.2-V59M bypass the regulation via ATP by increasing the channel open probability and reducing the ability of ATP to close the channel (*2*). This mutation-induced shift in the working range of K_ATP_ channels within PV-INs disturbs their normal feedback operation, impeding finely tuned high-frequency network activity such as SPWs and gamma oscillations (*27, 39, 40*), and permitting epileptic activity.

Our data indicate that dysfunctional PV-INs are crucial for the phenotype of patients with DEND syndrome. The similarity of effects induced by bath application of diazoxide and by PV-IN specific expression of the K_ir_6.2-V59M mutation suggests these fast-spiking interneurons with K_ATP_ channels play a dominant role in cognition-associated network activity. Our data also indicate that expressing K_ATP_ channels under control of the PV promoter provides a valid model of hippocampal K_ATP_ channel dysfunction in patients, in which the prevalence of mutated K_ATP_ channels in distinct cell types is controlled by the endogenous promotor.

Patients with DEND syndrome suffer from a range of neurological disabilities, including deficits in learning and memory, and difficulties in attention, perception, visuospatial abilities and sleep (*8, 9, 41*). Impairments in energy demanding network phenomena, such as SPWs and gamma oscillations, can potentially cause most of these higher brain functions (*16, 22, 27, 42*). Altered synaptic plasticity, contextual and spatial memory have previously mechanistically been linked to modified K_ATP_ channels in mice (*11, 43*). Our data demonstrate that enhanced K_ATP_ channel activity gives rise to impaired gamma oscillations and SPWs that underlie the neuropsychological problems. As gamma oscillations can be measured non-invasively and SPWs are with increasing frequency routinely measured in epilepsy patients, this makes them ideal correlates to further investigate cognitive deficits and treatment options in DEND syndrome patients.

Our *in vivo* data highlight another parameter that should be taken into consideration when characterizing DEND syndrome: circadian shifts in gamma activity. PV-V59M mice were unable to increase gamma power during wakefulness, a shift that may be expected to promote information perception and selection, behavioral adaptation and memory retrieval (*44*), all of which are affected in patients with DEND syndrome (*8, 9, 41*). Since locomotor activity was similar between mutant and control mice, our findings do not result from a lack of sensory stimuli or a general loss of circadian regulation in PV-V59M animals but from an inability to enhance gamma oscillations during wakefulness. Long-term EEG recordings are required to test whether a similar electrophysiological feature occurs in patients with DEND syndrome (*8, 41*).

Together with the increased susceptibility to epileptic activity in PV-V59M mice, which mirrors the phenotype of patients with DEND syndrome and is in line with the role of PV-INs in the control of epileptic activity (*45, 46*), our study identifies straightforward electrophysiological readouts for studying DEND syndrome in a mouse model: reduced SPW occurrence, altered gamma activity and increased seizure susceptibility. Our mouse model and experimental read-outs will prospectively help elucidate whether treatment of the neurological problems of patients with DEND syndrome with sulfonylureas was ineffective (*8, 11, 47–49*), because of unfavorable pharmacokinetics (*14*) or because of mutation-induced drug insensitivities (*2*). It will also facilitate testing the effectiveness of newly developed drugs targeting K_ir_6.2 channels (*50*). Hence, our study lays the foundation for better clinical diagnoses and new therapeutic strategies for DEND syndrome patients.

## Acknowledgements

We thank Gudrun Bethge for technical assistance and the members of the medical experimental center (MEZ), Leipzig University, for animal care. We thank Jan-Oliver Hollnagel for providing matlab scripts for data analysis of *in vitro* acquired SPWs and gamma oscillations. We thank Manfred Heckmann for fruitful discussions.

## Author contributions

M.E.B., J.K., J.E. and K.L. contributed in data acquisition, analysis, interpretation of the data and writing of the manuscript. J.V. created a new software tool for gamma resonance analysis. F.M.A. contributed to the conception of the work and the interpretation of the data. J.V. and F.M.A. contributed to the reviewing and editing of the manuscript.

## Additional information

Supplementary materials are available for this paper.

Correspondence should be addressed to

## Supplementary Materials

Materials and Methods

Figs. S1 to S2

References (*51-63*)

