## Supplement for "K_ATP_ channel mutation disrupts hippocampal network activity and nocturnal γ shifts"

†These authors contributed equally

*Corresponding author

#### Supplementary Materials

##### *Materials and Methods*

*Experimental license*

Animal experiments were approved by the regional authority of the State of Saxony (T 09/16, T 10/20, TVV 46/16, TVV 07/18, TVV 74/21) and performed in accordance with the guidelines of the European Communities Council.

*Mice*

For control and pharmacology experiments (Fig. 1) wildtype C57BL/6 mice were used. For all other experiments (Fig. 2-4/S1-2), genetically modified mutant mice expressing the $K_{ir}6.2$-V59M transgene selectively in $PV^+$-interneurons (PV-INs) were generated by using the Cre-loxP system, essentially as described previously (*12*). The $K_{ir}6.2$-V59M transgene, embedded in the ROSA26 locus, is preceded by a loxP-flanked stop sequence and followed by an FRT-flanked IRES GFP cassette. We generated PV-V59M mice by crossing mice heterozygously expressing this transgene (ROSA26StopKir6.2-V59M$^{lox/+}$ mice) with mice homozygously expressing Cre recombinase under the control of the parvalbumin (PV) promotor (B6;129P2-*Pvalb*$^{tm1(cre)Arbr}$/J, (*51*)). PV-V59M mice express a single copy of the Kir6.2-V59M transgene and two copies of the endogenous wild-type $K_{ir}6.2$ gene selectively in PV-INs. PV-tdTomato-V59M mice for cellular analyses (Fig. 3, S1) were then created by further crossing PV-V59M mice with the homozygous Ai14 mouse line (kindly provided by Hongkui Zeng, Allen Institute, Seattle, USA). These additionally express the fluorescent protein tdTomato (kindly provided by the laboratory of Roger Y Tsien, UC San Diego, USA) heterozygously in PV-INs enabling identification of PV-INs by live microscopy. Because $K_{ir}6.2$ requires SUR1 for surface expression, it is expected that expression of the $K_{ir}6.2$-V59M will not lead to overexpression of the $K_{ATP}$ channel (*12*, *52*, *53*). Littermates, heterozygously expressing PV-Cre recombinase (and heterozygous tdTomato for Fig. 3/S1) but not the $K_{ir}6.2$-V59M transgene, were used as control animals in all experiments (denoted as 'littermates' throughout the figures and text). Scientists conducting the experiments and analyses were blinded to the genotypes of the mice. Mice were maintained on a 12-hour light-dark cycle (6 a.m. to 6 p.m.) and had ad libitum access to water and food.

*Slice preparations and solutions*

For patch-clamp recordings, mice (p20-30) of either sex were deeply anesthetized using isoflurane (0.5 – 1%, Baxter, Unterschleißheim, Germany) and transcardially perfused with ice-cold *N*-Methyl-D-glutamine (NMDG)-containing slicing solution for about 1-2 min until the cardiovascular circulation was fully exchanged. Brains were quickly removed, glued with histoacryl tissue glue (B. Braun, Melsungen, Germany) onto a pre-cooled Teflon specimen holder and tilted to ~12° in fronto-occipital direction (*54*). Horizontal entorhinal-hippocampal slices of 300 µm thickness were prepared in ice-cold NMDG-slicing solution using a VT1200 S vibratome (Leica Biosystems, Wetzlar, Germany). Slices were then incubated in NMDG-slicing solution at 35 ± 2 °C for 12 min and stored in a HEPES-containing storage solution at room temperature until use. Recordings were performed in a submerged recording chamber with artificial cerebral spinal fluid (aCSF) prewarmed to 33 ± 2 °C using an inline-heater (SH-27B, Warner Instruments, Holliston, USA) at a flow rate of 5-8 ml/min. The NMDG-slicing solution contained (in mM): 92 NMDG, 20 HEPES, 30 $NaHCO_3$, 2.5 KCl, 10 $MgCl_2$, 0.5 $CaCl_2$, 1.2 $NaH_2PO_4$, 25 D-Glucose, 5 sodium ascorbate, 1 kynurenic acid, 5 *N*-acetyl-L-cysteine, 3 sodium pyruvate. The HEPESstoring solution contained (in mM): 92 NaCl, 20 HEPES, 30 $NaHCO_3$, 2.5 KCl, 1 $MgCl_2$, 2 $CaCl_2$, 1.2 $NaH_2PO_4$, 25 D-Glucose, 5 sodium ascorbate, 1 kynurenic acid, 5 *N*-acetyl-L-cysteine, 3 sodium pyruvate. Artificial CSF (aCSF, extracellular recording solution) contained (in mM): 129 NaCl, 1.25 $NaH_2PO_4$, 1.8 $MgSO_4$, 1.6 $CaCl_2$, 3 KCl, 10 D-Glucose, 21 $NaHCO_3$. All solutions were saturated with a mixture of 95% $O_2$ and 5% $CO_2$, adjusted to pH 7.3-7.4 and to an osmolarity of 305-315 mosmol/l (NMDG-slicing solution), or 300 ± 5 mosmol/l (HEPES-storage solution, aCSF). The glucose concentration (10 mM) was chosen to preserve slice viability and the capacity for generating high-frequency network oscillations (*55*).

Patch pipettes (2-6 MΩ) were pulled from filamented borosilicate glass (in mm: inner ∅ 1.16, outer ∅ 2.0, #1807533, Hilgenberg, Germany). For PV-IN whole-cell recordings the intracellular (pipette) solution contained (in mM): 150 K-gluconate, 3 ATP-Mg, 0.3 GTP-Na, 10 HEPES-K, 10 KCl, 0.05 EGTA (free acid), 5 GABA (liquid-junction (LJ) potential of 0 mV). For pyramidal cell (PC) whole-cell recordings the intracellular solution contained (in mM): 126 KCl, 4.5 ATP-$Na_2$, 0.3 GTP-Na, 10 HEPES, 1 NaCl, 10 EGTA (free acid), 4.5 $MgCl_2$, 1.25 $CaCl_2$ (LJ potential of 13 mV). For single-cell patch-clamp recordings (cf. Fig. 3A-H), 4 mM kynurenic acid and 10 μM GABAzine (SR-95531) were added to the aCSF to block ionotropic glutamatergic and GABAergic receptors. For paired recordings (cf. Fig. 3I-K), 40 μM CNQX and 50 μM D-AP5 were added to the aCSF to block NMDA- and AMPA-receptors.

For local field potential (LFP) recordings, mice of either sex were used. For recording spontaneously occurring SPWs, mice aged 7-8 weeks were used (Fig. 1D-E: 12 slices from 7 wild-type animals, (E) control n of SPWs=9746; diazoxide n=4760. Fig. 2C-D: littermates: 37 slices from 6 mice, (D) n=27412 SPWs. PV-V59M: 40 slices from 6 mice, (D) n=19848 SPWs). For recording gamma oscillations, mice aged 12-15 weeks were used (Fig. 1G-H: n=21 slices from 8 wild-type animals; Fig. 2G-H: 13 slices each from littermate and PV-V59M mice). Mice were decapitated under isoflurane anesthesia, the brain rapidly removed, and cut in ice-cold aCSF into 400 μm horizontal entorhinal-hippocampal slices, as described above. Slices were directly transferred to a storage and recording interface chamber and perfused with aCSF at 36°C for at least 2 hours (flow rate 1.8-2.0 ml/min) for slice recovery before the actual experiments were started. Extracellular borosilicate glass electrodes were pulled from filamented borosilicate glass (3 MΩ resistance; in mm: inner ∅ 1.16, outer ∅ 2.0, #1807533, Hilgenberg, Germany), filled with aCSF, were used for LFP recordings. For induction of gamma oscillations (Fig 1F-H, 2E-H), slices were superfused with 200 nM kainate dissolved in aCSF. $K_{ATP}$ channels were pharmacologically opened by applying 300 μM diazoxide (dissolved in 1 M NaOH with a volume fraction of 0.12 % in the final solution; Fig. 1). All substances were purchased from Sigma-Aldrich/Merck KGaA, Darmstadt, Germany, except for D-glucose and $NaHCO_3$ (Carl-Roth, Karlsruhe, Germany), and kainate (Tocris Bioscience, Bristol, UK).

*In vitro local field potential recordings and data acquisition*

Extracellular LFPs were recorded in the pyramidal layer of the hippocampal area cornu ammonis 1 (CA1). LFP signals were 100-fold pre-amplified using an EXT-10C amplifier, low pass filtered at 3 kHz (LPBF-01GX Filter) and 10-fold amplified (PA-2S amplifier, all by npi Electronic GmbH, Tamm, Germany). All signals were digitized at 10 kHz (1401 interface, CED, Cambridge, UK) and recorded using Spike2v.9 (Cambridge Electronic Design Ltd, Milton/Cambridge, UK).

*SPW detection and analysis*
For SPW detection, DC shifts in the LFP signal were first removed with a time constant of 0.5 s using Spike2. SPWs were then analyzed offline with MATLAB software (R2022a, The MathWorks, Inc., Natick, USA) using a custom-written script. Data were analyzed in 2 min bins. After applying a low pass filter to the data (cut frequency 80 Hz, 8$^{th}$ order, symmetrical Butterworth filter), the offset was corrected by subtracting the median of the low pass signal. A threshold crossing level (i.e., a multiple of the standard deviation crossing the mean of the low pass data), was used for detecting SPW events). These were identified, when the crossing level subsequently crossed twice (ascending and descending) the positive deflections of the low pass signal. Detection reliability was confirmed by eye and the threshold was adjusted globally for this recording, if necessary. The threshold was kept constant for pharmacological SPW experiments within one slice, for which two distinct time points were analyzed (baseline vs. diazoxide treatment, Fig. 1). For each data segment above the crossing level, the script searched for the local maximum, i.e., the SPW peak. This was then used to calculate the SPW amplitude and the number of SPWs per minute. For an exact determination of the SPW durations and amplitudes, a refinement of the threshold was performed for each SPW event by adding 15 ms of the low pass signal before (start) and after (end) the crossing levels. The median was calculated from these two 15 ms segments. If this median exceeded the crossing level between the start and three datapoints after the crossing level, the actual beginning of a SPW was redefined. The end of a SPW was redefined, if the median fell below the peak of the SPW in the range of 3 datapoints before the crossing level and the end of the SPW. The SPW duration was determined as the distance between the beginning and end point for each individual event. Events with durations below 5 ms or above 200 ms were discarded and adjacent events (inter-event interval < 15 ms) were merged. The SPW amplitude was calculated by subtracting the SPW peak, characterized as the maximum above the global baseline (i.e., the mean of the low pass filtered signal for each 2 min bin), from the local minimum of each SPW event. All SPW events were grouped (*56*, *57*) according to their treatment (control vs. diazoxide and littermates vs. PV-V59M, respectively).

*Gamma oscillation analysis*
Gamma oscillations were analyzed in MATLAB as described earlier (*58*, *59*). Briefly, LFP data segments consisting of the last 5 min of a 45 min long wash-in period were subdivided into sections of 30 s. Data were band pass filtered between 5-200 Hz using a fast Fourier transform (FFT) filter. Data were then processed using Welch´s algorithm and an FFT with a size of 8192 for generating the corresponding power spectral density (PSD) plots with a resolution of 1.2207 Hz. The frequency at the maximum of the PSD revealed the peak frequency. The integral of the PSD curve from 30-100 Hz relative to the integral of the total PSD curve (0.5-100 Hz) determined the relative gamma power. The median of the 30 s subdivisions was calculated for the peak frequency and relative gamma power.

*Patch-clamp electrophysiology from neurons in acute hippocampal slices*
Whole-cell patch-clamp recordings were conducted under visual guidance using an upright epifluorescence microscope (BX51WI, Olympus Corporation, Tokyo, Japan). The somata of PV-INs and PCs were located in the pyramidal layer of the CA1 subfield. PV-INs of PV-V59M mice were identified by the red fluorescence of the tdTomato fluorophore using LED excitation (pE-100, CoolLED, Andover, UK) and appropriate excitation/emission beam splitter. TdTomato was only expressed (under the PV-Cre promotor) for intracellular recordings depicted in Fig. 3 and S1.

After obtaining the whole-cell configuration, square pulses (1 s) with increasing amplitude were applied in current-clamp mode to identify neurons by means of their firing patterns. PV-INs showed fast-spiking non-adapting behavior, a short action potential (AP) half width and a prominent afterhyperpolarization, whereas PCs showed a decrease in AP amplitude and frequency during the depolarizing step, lower spiking frequencies and broader AP half widths. For data acquisition in all cellular recordings, EPC 10 USB DOUBLE Patch clamp amplifiers and the Patchmaster Software (Heka Elektronik GmbH, Reutlingen, Germany) were used (acquisition rate 100 kHz, filtered at 30 kHz). Current-clamp recordings were obtained in bridge mode with 100% series resistance ($R_s$) compensation. In voltage-clamp, $R_s$ was constantly monitored and automatically compensated up to 70% to keep the effective $R_s$ at a constant predefined value (typically 10-15 MΩ). Analysis of the patch-clamp electrophysiology was performed using custom-written scripts in the Igor Pro 8.04 software (WaveMetrics, Inc., Portland, USA).

*Single-cell patch-clamp recordings from PV-INs*

After establishing the whole-cell patch-clamp formation, PV-INs were voltage-clamped at -70 mV for at least 5 min until the membrane resistance ($R_m$) remained stable. PV-INs that showed continuous spontaneous firing or required a holding current exceeding -300 pA were discarded. $R_m$ was calculated from the steady-state current in response to a 5 mV hyperpolarization from the holding potential. The membrane potential ($V_m$) was measured in current-clamp mode with 0 pA holding current (fig. S1A-B: littermate n=35 PV-INs from 17 mice, PV-V59M n=36/20).

*Firing properties*

In current-clamp mode, ramp stimuli (1000 pA in 10 ms, fig. S1C: littermate n=15 PV-INs of 5 mice, PV-V59M n=17/7) were applied to determine the current needed to evoke a first action potential (AP threshold). A sinusoidal stimulus, linearly increasing in frequency (100-750 Hz within 8 s), was used to obtain the maximum AP frequency until the cell failed to fire (fig. S1D: littermate n=13 PV-INs from 5 mice, PV-V59M n=12/5).

*Intrinsic membrane potential oscillations*

In current-clamp mode, perithreshold intrinsic membrane potential oscillations were evoked by 2 s depolarizing current steps from -70 mV (0 to 600 pA in 50 pA steps; Fig. 3A-E: littermates n=10 PV-INs from 6 mice, PV-V59M n=15/9). Within this current step range, the most depolarizing step that evoked some APs but at least 1s without AP firing was repeated 5 to 10 times for each neuron and was used to calculate the power spectral density (PSD) of the oscillatory membrane activity. The neuronal membrane potentials at which the intrinsic membrane potential oscillations were measured were not different between groups ($V_m$: littermates: -53.79 [8.84]) mV vs. PV-V59M: -55.32 [7.01] mV, t-test p=0.334), which excludes voltage-dependent mechanisms for group differences in intrinsic oscillations. Gamma power was computed for each cell by integrating the area under the curve of the PSD from 30 to 100 Hz, respectively. The term "peak frequency" is used to refer to the peak of the mean PSD averaged from at least 5 sweeps.

*Gamma resonance*

Resonant properties of the cells were obtained by applying a subthreshold ZAP (impedance amplitude profile) stimulus, consisting of a pseudo-sinusoidal current of constant amplitude and linearly increasing frequency (10-100 Hz), for 10 s (Fig. 3F-H: littermates n=14 PV-INs of 8 mice; PV-V59M n=15/11). To find the AP threshold, single ZAP stimuli were applied (0 to 400 pA in

50 pA steps) until the cell started firing. From this AP threshold, the input current was decreased (in 10 pA steps) until the subthreshold range was reached, which was defined as the first step which failed to evoke APs. At this current step, the ZAP stimuli protocol was then repeated 4 to 8 times for each cell and membrane voltage waves were averaged for impedance analysis (subthreshold $V_m$: littermates: -67.73 [8.15]) mV vs. PV-V59M: -69.65 [10.90] mV, t-test p=0.996). Resonance analysis was performed, as previously described (*34*, *60*). In short, the curve of the impedance profile (Z(f)) was obtained from the ratio of the fast Fourier transform (FFT) of the output (voltage) and the input (current) waves (Z(f) = FFT[V(t)]/FFT[I(t)]). The impedance is a complex quantity ($Z(f) = Z_{Real} + iZ_{Imaginary}$) with $Z_{Real}$ being the resistive component and $Z_{Imaginary}$ the reactive component of the impedance. The complex impedance can be plotted as a vector for each given frequency with the magnitude provided by the following expression:

$$|Z|(f) = \sqrt{Z_{Real}^2(f) + Z_{Imaginary}^2(f)}$$

We use the term 'impedance' to refer to the magnitude of the impedance vector. For quantification of the gamma resonance, impedance data were fitted with a polynomial curve between 20 and 100 Hz to reveal the frequency at the peak of the impedance amplitude ($|Z_{max}|$), defined as the preferred frequency ($f_{Zmax}$) for each cell. Throughout the text, we refer to $|Z_{max}|$ as $Z_{max}$. Resonance strength was quantified by calculating the Q value as the ratio between $Z_{max}$ and the magnitude of the impedance at 20 Hz (referred to as $Z_{20Hz}$). Frequencies below 20 Hz were not plotted in the impedance profile graphs to avoid low frequency distortions.

*Paired recordings*

Paired recordings were obtained between PV-INs and PCs with their somata in the CA1 subfield. Both cells were identified by their distinct firing patterns. Until the paired recording was established, the presynaptic PV-IN was held in voltage-clamp at -70 mV with regular $R_s$ compensation. During the experiment, the PV-IN was held in current-clamp mode and the postsynaptic PC was held at -70 mV (corrected for the LJ potential of 13 mV) with automatic $R_s$ compensation in voltage-clamp mode. Presynaptic APs were evoked by applying brief (1 ms) current pulses with increasing amplitude until reliable APs occurred. In case of spontaneous firing, the PC was hyperpolarized (max. to -90 mV). To obtain the paired-pulse ratio (PPR) of a PV-IN-PC-pair, paired pulses were induced every 5 s with increasing inter-stimuli intervals (4, 10, 20, 50, 100 ms).

*Analysis of the IPSC amplitude, failure rate and PPR*

In the case of spontaneous firing of either the PV-IN or the PC, the affected parts of the recording were discarded. The first IPSCs of the PPR protocol were used to determine the IPSC amplitude, kinetics and failure rate. The peak IPSC amplitude was measured as the difference between the minimum of the IPSC and the average of a 1 ms baseline period just prior the beginning of the IPSC. In order to analyze IPSC amplitudes measured at different PC holding potentials (Fig. 3J: littermates n=10 PV-IN-PC pairs of 9 mice; PV-V59M n=11/10), we corrected for the reversal potential of $GABA_A$ receptors, which was assumed to be 0 mV with our intracellular and extracellular solutions. Failures of synaptic transmission were defined as the lack of a post-synaptic response between 1 and 3 ms after a pre-synaptic AP, and were visually detected by a person blinded to the genotype of the mouse. Data from neuronal pairs with a failure rate of ≥ 25%

were discarded (littermates n=2 pairs in 2 mice; PV-V59M n=3/3) following (*61*), who found a mean failure rate of 3 ± 2% for the CA1 PV-IN-PC-synapse. Due to a high variability in the post-synaptic responses, we calculated the PPR as the ratio of the mean of all second IPSC amplitudes (per ISI) to the mean of all first IPSC amplitudes (of all ISIs) for each pair (Fig. 3K: littermates n=7-10 pairs in 6-9 mice; PV-V59M n=9-11/9-10).

*Stereotactic intrahippocampal electrode implantation, in vivo data acquisition and analysis*

Intrahippocampal *in vivo* LFP recordings were performed, as previously described (*62*). Briefly, the telemetric system from Data Science International (DSI, St. Paul, MN, USA) was used with mouse-implantable ETA-F10 transmitters for continuously measuring LFPs and activity of freely-moving mice. For electrode implantation, mice (8-12 weeks old, n=8 littermate and n=9 PV-V59M mice, respectively) received preemptive metamizole analgesia (200 mg/kg bodyweight (b.w.)), were deeply anesthetized using a combination of a single dose ketamine (30 mg/kg b.w.) and continuous administration of 0.25-3% isoflurane (in 100% oxygen), and received local analgesia by use of a lidocaine gel on the head skin. Mice were head-fixed in a stereotactic frame and after opening the skin of the head, three holes were drilled in the skull. A sml-coated tungsten wire (ø = 50 µm, WireTronic Inc., Volcano, CA, USA) was inserted with its uninsulated end into the coiled recording electrode of the transmitter. This connection was mechanically compressed and subsequently insulated with dental cement (Unifast TRAD, GC America Inc., Alsip, IL, USA). The other, end (insulated except for the tip) of the wire was then inserted in the CA1 pyramidal layer of the right hippocampus (0.150 mm lateral, 0.182 mm posterior from bregma to 0.121 mm ventral below cortical surface, The Mouse Brain, Paxinos & Franklin) as the recording electrode. The outer part of the inserted electrode was laterally stabilized with a screw (3/32-inch, stainless steel, J.I. Morris, Oxford, MA, USA) and dental cement. The second, coiled reference electrode was uncoiled and twisted around a screw that was screwed into the skull as an epidural reference above the left cerebellar cortex. The two electrodes redirect their electrical signals through a DSI transmitter, which was implanted subcutaneously on either side of the thorax. Mice received 1 ml of subcutaneously injected saline as fluid substitution. Post-operative analgesia was applied by 300 mg metamizole/100 ml drinking water, with 4 ml of 20% glucose added. Mice were checked daily for health and bodyweight. In case of pain, subcutaneous metamizole (200 mg/kg b.w.) was given: this was sometimes necessary within the first two days post-surgery. Data were sampled at 1 kHz and transmitted over a frequency band of 0.5-200 Hz telemetrically from the implanted DSI transmitter to the receiver boards, which recorded the signal using Ponemah software (v6.5) from DSI. Movement activity counts were measured as a relative change of signal strength dependent on both distance and speed of animal movement. Activity counts were sampled and transmitted to the receiver board with a frequency of 1 Hz.

*Seizure, gamma and movement activity analysis*

Data analysis was performed using NeuroScore v3.0 from DSI. To eliminate postoperative influences, data was analysed on postsurgical days 4-6 for each animal. To identify seizures, the automated threshold detection of Neuroscore was used. The threshold was set for each recording on the basis of baseline LFP activity so that over-baseline events were adequately detected. All events were reviewed by a blinded person with the help of video recordings if applicable. Only events with a sufficiently high amplitude, typical spiking pattern, no movement artifacts and a minimum duration of 5 s were classified as seizures. Identified seizures were reviewed by another person, blinded to genotype, for confirmation. The event durations were manually measured using Neuroscore software.

For network oscillation analysis, the power of the sampled data was computed as the integral of the power spectrum for a given frequency range based on an FFT. Data were computed with a bin size of 2 s and a sliding average of 50% overlap. The relative power was computed as the ratio between the power of the gamma band (30-100 Hz) and the total power of the signal (0.5-200 Hz). For statistical analysis, the relative power of gamma oscillations (30-100 Hz) was averaged for day (6 a.m. to 6 p.m.) and night (6 p.m. to 6 a.m.) according to the day and night rhythm of the animal housing. Movement activity counts were accumulated hourly for three consecutive days (postoperative days 4-6) and averaged for each animal before averaging the group.

*Cresyl violet staining and electrode positions*

To verify the tungsten electrode positions in the hippocampal area CA1, mice were deeply anesthetized using combined ketamine/xylazine anesthesia (100 mg/kg b.w. (WDT, Garbsen Germany) and 5 mg/kg b.w. (Bayer, Leverkusen, Germany), respectively) and transcardially perfused with 4% of paraformaldehyde (PFA). Brains were removed, fixed in PFA overnight and incubated in 30% sucrose for another night. The part of the brain harbouring the area of interest was embedded in a Tissue Tec CryoMold within O.C.T. compound solution (Sakura Finetek Europe B.V., Alphen aan den Rijn, NL) and subsequently frozen in 2-methylbutane (Carl Roth, Karlsruhe, Germany) using liquid nitrogen. Frozen tissue was cryosectioned (30 µm) with a cryotome (CM3050 S, Leica Biosystems, Nussloch, Germany) and dried on a microscope slide. Slices were stained according to the cresyl violet staining procedure of FD neurotechnology (FD Cresyl Violet Solution User Manual™, PS102v.2011-1), comprising incubation of slides in xylene, ethanol, cresyl violet (Sigma-Aldrich, 1% for 6 min) and glacial acetic acid. Slices were embedded on the microscope slide using mounting medium (Immunoselect Antifading Mounting Medium, dianova, Hamburg, Germany).

*Immunohistochemistry of PV-INs*

A PV-tdTomato mouse (age p24) was transcardially perfused with 4% PFA (as described above) and 40 µm thick sagittal hippocampal slices were cut using a vibratome (VT1000S, Leica Biosystems). After washing slices in PBS, slices were blocked and permeabilized with a mixture of PBS, fetal calf serum and triton-x 100. Slices were incubated overnight with the primary anti-RFP antibody (polyclonal, originated from rabbit, Rockland Immunochemicals, Inc., Pottstown, PA, USA, #600-401-379) with a concentration of 1:200. Slices were then washed with PBS and incubated for two hours with the Cy3 donkey-anti-rabbit secondary antibody (1:200 dissolved in 50% glycerol, Jackson ImmunoResearch, West Grove, PA, USA, #711-165-152). Slices were embedded in mounting medium on an objective slide. High-resolution images were taken of the hippocampal formation using a confocal microscope (Olympus IX71/Fluoview, 533 nm excitation, 580/60 nm emission filter). Images were stitched using an ImageJ plugin (*63*).

*Statistical analyses*

For statistical comparison, data were either presented as probability density functions (PDF), as normalized cumulative density functions (CDF), or as single data points with corresponding boxplots that displayed the median and the range of the $25^{th}$ to $75^{th}$ quartiles and whiskers denoting the last data points within quartiles + 1.5* interquartile range (IQR). Data were described as median and IQR in the text, if not otherwise stated. Statistical comparison of histograms (PDFs, CDFs) was conducted using the Kolmogorov-Smirnov test. For normally distributed numerical data and non-parametric data, the two-tailed student's t-test (t-test) and the Mann-Whitney-U test (MW) were used, respectively (IgorPro, Wavemetrics, Lake Oswego, OR, USA). To compare dependent data, either the paired t-test or the Wilcoxon signed-rank test (WSRT) were used, according to the distribution of the data. The two-tailed Fisher´s exact tests (Fisher´s) were used to compare categorical data (GraphPad Software, San Diego, CA, USA). P-values (p) < 0.05 were considered as statistically significant, whereby single asterisks denote p-values < 0.05, double asterisks < 0.01 and triple asterisks < 0.001.

*Data and code availability*

Data of this study are available from the corresponding author upon request. Codes that were used for analyzing data of this study are available at https://github.com/KristinaLippmann.

***Supplementary figures***

Figure S1

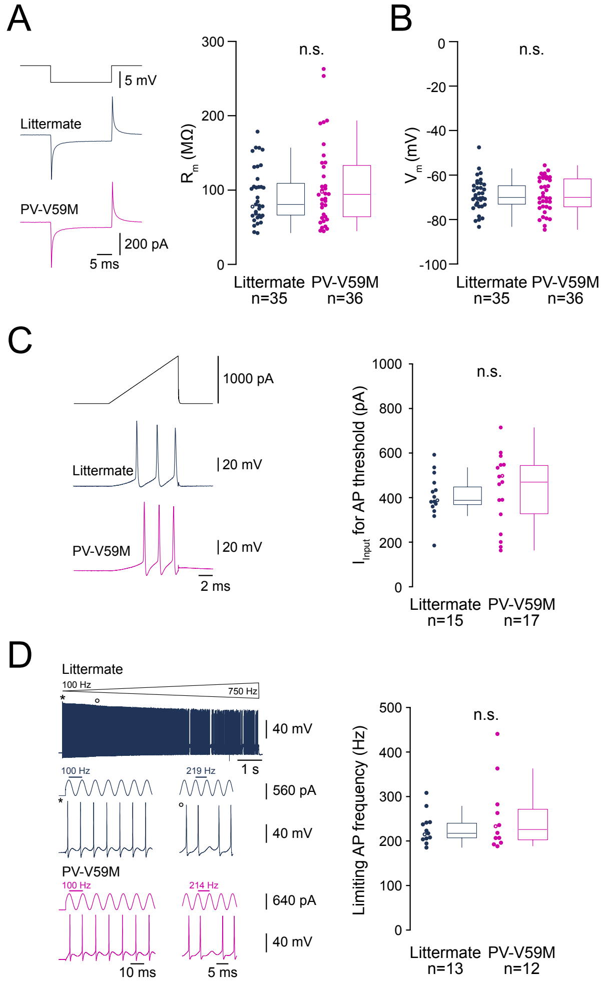


**Supplementary fig. 1, relating to Fig. 3. Basic intrinsic properties of PV-INs are unchanged in PV-V59M mice.** Open circles depict examples shown on the left side of the panels. **A.** Left: Membrane resistance ($R_m$) calculated from steady-state current responses (middle and lower trace) to a hyperpolarizing voltage step (top trace). Right: $R_m$ for PV-INs from littermates (80.7 [44.3] MΩ) and PV-V59M mice (94.2 [70.6] MΩ, median [IQR]). **B.** Resting membrane potential ($V_m$) of PV-INs from littermates (-70.1 [8.9] mV) and PV-V59M mice (-70.0 [13.1] mV). **C.** Left: The minimal depolarizing current ($I_{Input}$) that induced one or more APs, determined by applying a ramp stimulation (top trace, 0 to 1000 pA over 10 ms). Middle and lower left panels show typical voltage responses. Right: Threshold currents for PV-INs from littermates (388 [84] pA) and PV-V59M mice (470 [221] pA). **D.** Left: Maximum AP frequency was determined by applying a linearly increasing sinewave from 100 to 750 Hz, displayed as cone, with a representative littermate $V_m$ (top trace). The middle and lower traces show representative segments at the starting frequency (100 Hz) and the frequency at which a first AP failure occurred. Littermates in blue, PV-V59M in magenta. Right: Highest failure-free frequency (littermates: 217 [36] Hz vs. PV-V59M: 226 [71] Hz).

### Figure S2

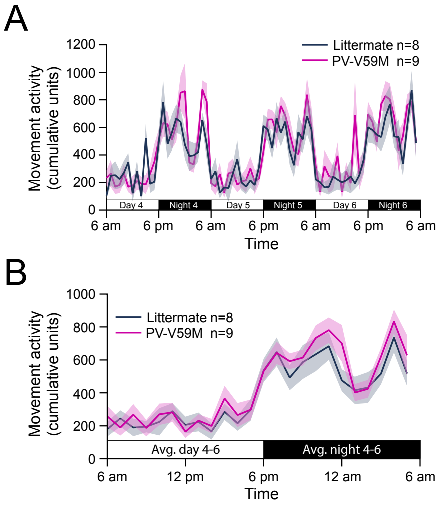


**Supplementary fig. 2, relating to Fig. 4. Movement activity in PV-V59M mice is unchanged from littermates. A.** Movement activity data grouped for littermates and PV-V59M mice (mean ± SEM) from post-operative days 4-6 and presented in cumulative NeuroScore units. **B.** Activity data from A, averaged over three days, displayed over one day and night cycle.
